# *Lelliottia amnigena* recovered from the lung of a harbour porpoise, and comparative analyses with *Lelliottia* spp.

**DOI:** 10.1101/2023.08.09.552678

**Authors:** David Negus, Geoffrey Foster, Lesley Hoyles

## Abstract

Strain M1325/93/1 (= GFKo1) of *Lelliottia amnigena* was isolated from the lung of a harbour porpoise in 1993. The genome sequence and antimicrobial resistance profile (genomic, phenotypic) of the strain were generated, with the genomic data compared with those from closely related bacteria. We demonstrate the recently described chromosomally-encoded AmpC β-lactamase *bla*_LAQ_ is a core gene of *L. amnigena*, and suggest new variants of this class of lactamase are encoded by other members of the genus *Lelliottia*. Although presence of *bla*_LAQ_ is ubiquitous across the currently sequenced members of *L. amnigena*, we highlight that strain GFKo1 is sensitive to ampicillin and cephalosporins. These data suggest *bla*_LAQ_ may act as a useful genetic marker for identification of *L. amnigena* strains, but its presence may not correlate with expected phenotypic resistances. Further studies are required to determine the regulatory mechanisms of *bla*_LAQ_ in *L. amnigena*.

## INTRODUCTION

*Lelliottia* spp. are Gram-negative, facultatively anaerobic bacteria of the family *Enterobacteriaceae*. The genus *Lelliottia* was created to accommodate species distinct from *Enterobacter sensu lato* based on *gyrB, rpoB, infB* and *atpD* gene sequence analyses, and comprises four species with validly published names (*Lelliottia amnigena, Lelliottia aquatilis, Lelliottia jeotgali* and *Lelliottia nimipressuralis*) and one with a non-valid name (“*Lelliottia steviae*”) (Brady *et al*. 2013; Kämpfer *et al*. 2018; Yuk *et al*. 2018; Lin *et al*. 2022). *Lelliottia aquatilis* represents a later heterotypic synonym of *L. jeotgali*, based on average nucleotide identity (ANI) and *in silico* DNA–DNA hybridization analyses (Wu and Zong 2019).

*Lelliottia* spp. have been associated with the commensal microbiota of flies and the Asian tiger mosquito (Guégan *et al*. 2020; Wiktorczyk-Kapischke *et al*. 2022), and isolated from fresh and waste water, soil, plants, air samples and fish (Heinle *et al*. 2018; Kämpfer *et al*. 2018; Yuk *et al*. 2018; Salgueiro *et al*. 2020; Reitter, Neuhaus and Hügler 2021; Leister and Hügler 2022; Thakur and Gauba 2022; Tran *et al*. 2022; Bilous *et al*. 2023; Suescun-Sepulveda, Rondón González and Fuentes Lorenzo 2023). Interest in *L. amnigena* is increasing as this bacterium has been associated with soft rot of economically important plant crops such as onion and potato (Osei *et al*. 2022). Only rarely have *L. amnigena* and *L. nimipressuralis* been associated with opportunistic disease in humans (Leal-Negredo *et al*. 2017; Martín Guerra, Martín Asenjo and Dueñas Gutiérrez 2018; Choi *et al*. 2021; Legese *et al*. 2022). There are few reports in the literature of the carriage of antimicrobial resistance (AMR) genes by *Lelliottia* spp., though a new chromosomally-encoded AmpC β-lactamase, *bla*_LAQ-1_, conferring resistance to ampicillin and several cephalosporins was recently described for an *L. amnigena* strain isolated from animal farm sewage in China (Li *et al*. 2022; El Zowalaty *et al*. 2023).

As part of a study of veterinary isolates thought to belong to the *Klebsiella oxytoca* complex (Smith-Zaitlik *et al*. 2022), we identified several atypical strains that were shown by *rpoB* gene sequence analysis to represent a range of different *Enterobacteriaceae* (Smith-Zaitlik 2021). Here, we report on one such strain recovered from the lung of a harbour porpoise (*Phocoena phocoena*). Using genome sequence data and comparative analyses, we demonstrate this is a strain of *L. amnigena* and compare its AMR gene profile with those of publicly available sequence data for the species.

## MATERIALS AND METHODS

### Isolation and phenotypic characterization of strain

Strain M1325/93/1 (herein referred to by our laboratory identifier, GFKo1) was isolated on Columbia sheep blood agar (Oxoid, Basingstoke, UK) from the lung of a harbour porpoise that was found stranded at Buckie on the southern coastline of the Moray Firth, north-east Scotland in June 1993. Tentative identification and biochemical characterization of the strain were made using the API 20E (bioMérieux) strip according to the manufacturer’s instructions under aerobic conditions at 37 °C. The isolate was also identified by matrix-assisted laser desorption-ionisation time-of-flight mass spectroscopy (MALDI-TOF) using the Bruker Microflex™ LT/SH MALDI-TOF MS Biotyper™. Antimicrobial sensitivity testing was performed by disc diffusion assays following guidelines from the European Committee on Antimicrobial Susceptibility Testing (EUCAST) v 13.1 for *Enterobacterales. Escherichia coli* ATCC 25922 was used as the reference strain for quality control purposes. All antibiotics were purchased from Oxoid, UK.

### DNA extraction and sequencing

DNA was extracted from an overnight culture (aerobic, 37 °C) of strain GFKo1 grown in nutrient broth (Oxoid) using the Qiagen DNeasy Blood and Tissue Kit (Qiagen). Extracted DNA was adjusted to a concentration of 0.2 ng/μL and treated using the Nextera XT DNA library preparation kit (Illumina) to produce fragments of approximately 500 bp. Fragmented and indexed samples were run on the sequencer using the MiSeq Reagent Kit v2 (Illumina; 250□bp paired-end reads) following Illumina’s recommended denaturation and loading procedures.

### Genome assembly and gene annotation

Raw sequence data were checked using fastqc v0.11.4 (https://www.bioinformatics.babraham.ac.uk/projects/fastqc/); no adapter trimming was required, and reads had an average Phred score >25. Genome data for strain GFKo1 were assembled using Megahit v1.2.9 (options: --min-contig-len 500 -r), with only contigs ≥500 nt in length retained. CheckM2 v0.1.3 (Chklovski *et al*. 2022) was used to determine the completeness and contamination of the genome sequence. Bakta v1.4.2 (database 3.1) (Schwengers *et al*. 2021) was used to annotate predicted genes within the genome.

### Accurate identification of genomes

Ribosomal multi-locus sequence typing (rMLST; (Jolley *et al*. 2012)) was used to identify the closest relative of strain GFKo1. OAT:OrthoANI v0.93.1 (Lee *et al*. 2016) was used to determine ANI values for the genome with publicly available *L. amnigena* genomes and type strains of closest relatives. Identities of publicly available genome sequences of *L. amnigena* (downloaded from NCBI GenBank on 19 March 2023; **Table 1**) were confirmed by comparison (OAT:OrthoANI) with the genome sequences of the type strains of the genus. These genomes were checked, annotated and identified as described above. Sourmash v4.6.1 was used to generate 31-kmer signatures for genomes, which were compared with one another to determine how similar genomes were to one another, and to identify genomes belonging to *L. amnigena sensu stricto* (Brown and Irber 2016). PhyloPhlAn3 (--diversity medium) was used to confirm the affiliation of all genomes with the genus *Lelliottia*.

**Table 1.** Sequence summary statistics for Bakta-annotated genomes included in this study.

| Strain | Accession | Source | Size (bp) | Contigs | GC content (%) | N50 | CDS | CheckM2 |  |
| --- | --- | --- | --- | --- | --- | --- | --- | --- | --- |
|  |  |  |  |  |  |  |  | Completeness (%) | Contamination (%) |
| M1325/93/1 (=GFKo1) | JAUBKL000000000 | Porpoise lung, UK | 4294992 | 200 | 53.1 | 46243 | 3954 | 100 | 0.06 |
| 155047<sup>T</sup> | GCA_022171985 | Human sputum, China | 4990088 | 98 | 53.7 | 358667 | 4707 | 100 | 0.20 |
| NCTC 12124<sup>T</sup> | GCA_900635465 | Soil | 4471442 | 1 | 52.9 | 4471442 | 4572 | 100 | 0.23 |
| 6331-17<sup>T</sup> | GCA_002923025 | Water, Germany | 4774414 | 37 | 54.2 | 202682 | 4474 | 100 | 0.00 |
| CCUG 25894<sup>T</sup> | GCA_004115925 | Elm tree, USA | 4616251 | 67 | 54.8 | 236780 | 4293 | 100 | 0.05 |
| PFL01<sup>T</sup> | GCA_002271215 | Jogaejeotgal, South Korea | 4603334 | 1 | 54.2 | 4603334 | 4237 | 100 | 0.01 |
| LST-1 | CP063663 | *Stevia*, China | 3576481 | 1 | 41.1 | 3576481 | 3187 | 100 | 0.03 |
| JCM 17292<sup>T</sup> | GCA_001550155 | Sediment, Arabian Sea | 4459111 | 26 | 40.9 | 658688 | 4004 | 100 | 0.19 |
| 2017H1G6 | GCA_004331765 | Soil, Denmark | 4606148 | 90 | 52.7 | 134684 | 4343 | 100 | 0.01 |
| 4928STDY7071390 | GCA_902160115 | Human faeces, UK | 4467891 | 28 | 55.3 | 476430 | 4119 | 100 | 0.00 |
| A167 | GCA_021498285 | Soil, Netherlands | 4662149 | 2 | 52.8 | 4520659 | 4344 | 100 | 0.05 |
| ENT01 | GCA_025641975 | Soil, USA | 4716124 | 59 | 52.9 | 212085 | 4402 | 100 | 1.32 |
| ERR1430553* | GCA_938039995 | Human faeces, China | 4361353 | 909 | 53.0 | 5972 | 4272 | 90.45 | 4.58 |
| ERR1430553* | GCA_905202905 | Human faeces, China | 3854042 | 799 | 53.4 | 5991 | 3704 | 88.98 | 5.15 |
| ERR5094855* | GCA_947072025 | Rainbow trout gut, France | 4359307 | 65 | 52.9 | 139247 | 4050 | 99.37 | 0.65 |
| FDAARGOS 1444 | GCA_019047465 | Unknown | 4505532 | 1 | 52.8 | 4505532 | 4169 | 100 | 0.15 |
| FDAARGOS 1446 | GCA_019048185 | Unknown | 4914411 | 5 | 52.6 | 4591698 | 4772 | 100 | 1.27 |
| FDAARGOS_1445 | GCA_019355955 | Unknown | 4599109 | 2 | 52.8 | 4504790 | 4287 | 100 | 0.06 |
| FDAARGOS_395 | GCA_002393405 | Soil, USA | 4469608 | 1 | 52.9 | 4469608 | 4130 | 100 | 0.01 |
| INSAq176 | GCA_021441185 | Fish, Portugal | 4422149 | 193 | 53.2 | 58074 | 4147 | 95.84 | 0.07 |
| JUb66 | GCA_003752235 | Unknown | 4572787 | 1 | 52.9 | 4572787 | 4205 | 100 | 0.02 |
| P13 | GCA_023970615 | Pig (sewage), China | 4622385 | 2 | 52.9 | 4555627 | 4316 | 100 | 0.90 |
| PTJIT1005 | GCA_022352085 | Water, India | 4550713 | 71 | 52.9 | 298940 | 4250 | 100 | 0.08 |
| TZW12 | GCA_016771075 | Water, Germany | 4694183 | 26 | 52.5 | 415957 | 4420 | 100 | 0.00 |
| TZW13 | GCA_016770995 | Water, Germany | 4830285 | 26 | 52.5 | 337333 | 4622 | 100 | 0.05 |
| TZW14 | GCA_016770935 | Water, Germany | 4516381 | 17 | 52.8 | 731232 | 4206 | 100 | 0.01 |
| TZW15 | GCA_016770975 | Water, Germany | 4756711 | 36 | 52.6 | 346396 | 4485 | 100 | 0.03 |
| TZW16 | GCA_016770955 | Water, Germany | 4756331 | 35 | 52.6 | 346396 | 4481 | 100 | 0.03 |
| UMA3121 | GCA_013337605 | Forest soil, Portugal | 4420612 | 19 | 52.9 | 559149 | 4091 | 100 | 0.00 |
| ZB04 | GCA_001652505 | Midgut of silkworm, China | 4616122 | 1 | 54.3 | 4616122 | 4205 | 100 | 0.03 |

\*MAGs; full names ERR1430553\_bin.131\_CONCOCT\_v1.1\_MAG, ERR1430553-bin.48 and ERR5094855\_bin.4\_metaWRAP\_v1.3\_MAG.

### Identification of AMR genes predicted to be encoded in genomes

Initially, the Resistance Gene Identifier [RGI 6.0.1, CARD 3.2.6; (Alcock *et al*. 2020)] was used to derive information on AMR genes predicted to be encoded in the genome of strain GFKo1. The genome sequence of GFKo1 was also searched for the allele of the chromosomal class C β-lactamase *bla*_LAQ-1_ (nucleotide accession MZ497396; (Li *et al*. 2022)) using Geneious Prime v2023.0.1. Based on the result of the *bla*_LAQ-1_ search, AMRFinderPlus v3.11.4 (database version 2023-02-23.1) (Feldgarden *et al*. 2022) and Bakta annotations were subsequently used for surveying AMR genes in genomes.

A BLASTP database was created using the amino acid sequence of MZ497396. Bakta-annotated protein sequences for all genomes (**Table 1**) were searched against this sequence, with hits >70 % coverage and >70 % identity retained. The ‘hit’ protein sequences were extracted from the .faa Bakta-annotated files using Biostrings v2.64.0 and used to create a multiple-sequence alignment (Clustal Omega v1.2.2; Geneious Prime v2023.0.1) with the protein sequences of the 12 AmpC β-lactamases (ACT-12, ACT-22, BIL-1, CMY-2, CMY-20, LAT-1, CFE-1, YRC-1, MIR-1, MIR-23, ACT-6, ACT-10) included in the study in which the functionality of the *bla*_LAQ-1_ protein was demonstrated (Li *et al*. 2022). A phylogenetic tree was created from the sequence alignment using PhyML v3.3.20180621 (Blosum62 matrix) (Guindon *et al*. 2010), with bootstrap values determined based on 100 replications. The tree was visualized using iToL v6 (Letunic and Bork 2019) with additional annotations made using Adobe Illustrator.

## RESULTS

### Characteristics of genome of GFKo1

Strain GFKo1 was recovered from the lung of a harbour porpoise that stranded in 1993. Although originally thought to represent a strain of *K. oxytoca, rpoB* gene sequence analysis done in the laboratory at Nottingham Trent University showed the strain was a representative of *L. amnigena* (Smith-Zaitlik 2021). This identification was supported by API 20E data (read after 24 and 48 h; code 1305173: *Enterobacter amnigenus* 1 90.4 %) and by MALDI-TOF with scores that reached 2.48, significantly above the 2.0 cut-off for species identification.

As *L. amnigena* has not previously been associated with marine mammals and there are few genome sequences available for the species, we generated the draft genome sequence of strain GFKo1 (20× coverage). The genome comprised 4,294,992 bp across 200 contigs (N50 46,243), and was predicted to encode 3,954 coding sequences, 80 tRNA, 1 tmRNA and 6 ribosomal RNA genes (**Table 1**). This information, together with its high completeness and low contamination (**Table 1**), demonstrated GFKo1’s genome was of high quality (Bowers *et al*. 2017).

rMLST (Jolley *et al*. 2012; Jolley, Bray and Maiden 2018) identified GFKo1 as *L. amnigena* (100 % identity). This is a rapid method that indexes variation of the 53 genes encoding bacterial ribosome protein subunits to integrate microbial taxonomy and typing. ANI analysis of GFKo1’s genome against the genomes of type strains of the genus *Lelliottia* confirmed GFKo1 as a strain of *L. amnigena*, sharing 98.31 % ANI with the type strain (NCTC 12124^T^, assembly accession GCA_900635465) of the species (Chun *et al*. 2018) (**Fig. 1a**).

**Fig. 1.**
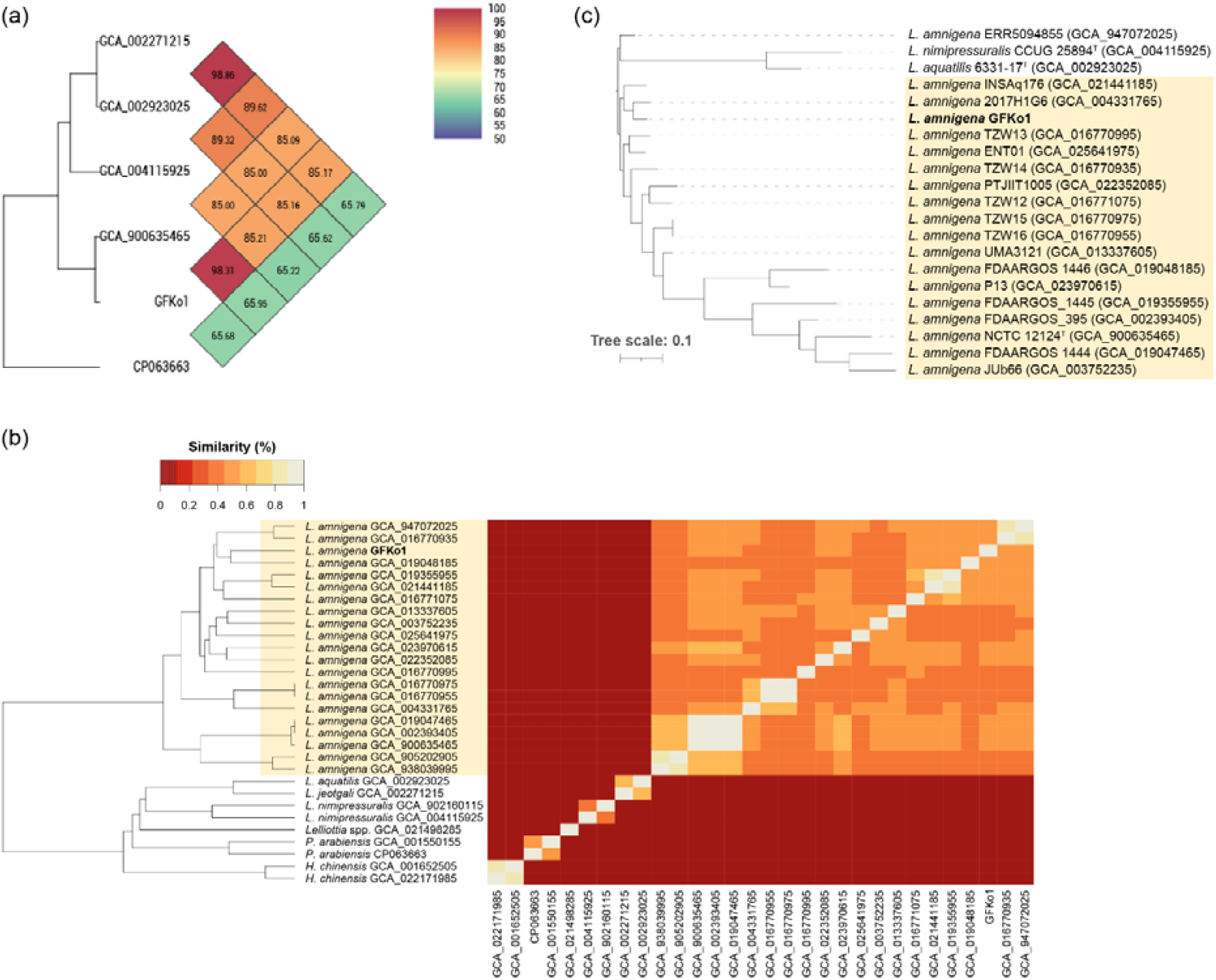
Strain GFKo1 is a representative of *L. amnigena*. (a) Heatmap generated by OAT:OrthoANI showing the ANI between GFKo1 and strains listed as type strains of *Lelliottia* species with valid and non-valid names. GFKo1 shares highest ANI (%) with the type strain of *L. amnigena* (accession assembly GCA_900635465). (b) Heatmap with unidirectional clustering showing the similarity of sourmash signatures across all genomes included in this study. The lighter the colour of the block on the heatmap, the more similar the two corresponding genome signatures. (c) RAXmL (best tree) generated by PhyloPhlAn3 from the proteomes of high-quality (>90 % completeness, <5 % contamination; **Table 1**) genome sequence data for the genus *Lelliottia*. The tree was rooted on the clade containing *L. nimipressuralis* and *L. aquatilis*. Scale bar, average number of amino acid substitutions per position. (b, c) The clade highlighted in light yellow represents *L. amnigena sensu stricto*.

### Curation of *Lelliottia* genome dataset

We downloaded the GenBank genome assemblies of all *Lelliottia* type strains (*n*=5) and all *L. amnigena* (*n*=22, excluding *L. amnigena* type) strains from NCBI GenBank (**Table 2**). All were checked for completeness and contamination using CheckM2 (**Table 1**). Except for metagenome-assembled genome (MAG) ERR1430553, all were of high quality (<5 % contamination, >90 % complete) (Bowers *et al*. 2017).

**Table 2.** Species identities of genomes included in this study as determined using different methods.

| Strain | Accession | NCBI ID | rMLST ID, % support | ANI with type strain genome |
| --- | --- | --- | --- | --- |
| M1325/93/1 (=GFKo1) | JAUBKL000000000 | *L. amnigena* | *L. amnigena* 100 % | *L. amnigena* 98.31 % |
| 155047<sup>T</sup> | GCA_022171985 | *Huaxiibacter chinensis* | *H. chinensis* 100 % | *H. chinensis* 100 % |
| NCTC 12124<sup>T</sup> | GCA_900635465 | *L. amnigena* | *L. amnigena* 100 % | *L. amnigena* 100 % |
| 6331-17<sup>T</sup> | GCA_002923025 | *L. aquatilis* | *L. aquatilis* 100 % | *L. aquatilis* 100 % |
| CCUG 25894<sup>T</sup> | GCA_004115925 | *L. nimipressuralis* | *L. nimipressuralis* 100 % | *L. nimipressuralis* 100 % |
| PFL01<sup>T</sup> | GCA_002271215 | *L. jeotgali* | *L. aquatilis* 90 % | *L. jeotgali* 100 % |
| LST-1 | CP063663 | "*L. steviae*" | *P. arabiensis* 100 % | *P. arabiensis* 99.13 % |
| JCM 17292<sup>T</sup> | GCA_001550155 | *P. arabiensis* | *P. arabiensis* 100 % | *P. arabiensis* 100 % |
| 2017H1G6 | GCA_004331765 | *L. amnigena* | *L. amnigena* 100 % | *L. amnigena* 98.41 % |
| 4928STDY7071390 | GCA_902160115 | *L. amnigena* | *L. nimipressuralis* 93 % | *L. nimipressuralis* 98.15 % |
| A167 | GCA_021498285 | *L. amnigena* | *L. amnigena* 100 % | *L. amnigena* 93.65 % |
| ENT01 | GCA_025641975 | *L. amnigena* | *L. amnigena* 100 % | *L. amnigena* 98.29 % |
| ERR1430553* | GCA_938039995 | *L. amnigena* | *L. amnigena* 54 % | *L. amnigena* 99.15 % |
| ERR1430553* | GCA_905202905 | *L. amnigena* | *L. amnigena* 57 % | *L. amnigena* 99.20 % |
| ERR5094855* | GCA_947072025 | *L. amnigena* | *L. amnigena* 100 % | *L. amnigena* 98.32 % |
| FDAARGOS 1444 | GCA_019047465 | *L. amnigena* | *L. amnigena* 100 % | *L. amnigena* 99.97 % |
| FDAARGOS 1446 | GCA_019048185 | *L. amnigena* | *L. amnigena* 100 % | *L. amnigena* 98.32 % |
| FDAARGOS_1445 | GCA_019355955 | *L. amnigena* | *L. amnigena* 100 % | *L. amnigena* 98.45 % |
| FDAARGOS_395 | GCA_002393405 | *L. amnigena* | *L. amnigena* 100 % | *L. amnigena* 99.97 % |
| INSAq176 | GCA_021441185 | *L. amnigena* | *L. amnigena* 100 % | *L. amnigena* 98.42 % |
| JUb66 | GCA_003752235 | *L. amnigena* | *L. amnigena* 100 % | *L. amnigena* 98.40 % |
| P13 | GCA_023970615 | *L. amnigena* | *L. amnigena* 100 % | *L. amnigena* 98.87 % |
| PTJIT1005 | GCA_022352085 | *L. amnigena* | *L. amnigena* 100 % | *L. amnigena* 98.85 % |
| TZW12 | GCA_016771075 | *L. amnigena* | *L. amnigena* 100 % | *L. amnigena* 98.45 % |
| TZW13 | GCA_016770995 | *L. amnigena* | *L. amnigena* 100 % | *L. amnigena* 98.30 % |
| TZW14 | GCA_016770935 | *L. amnigena* | *L. amnigena* 100 % | *L. amnigena* 98.24 % |
| TZW15 | GCA_016770975 | *L. amnigena* | *L. amnigena* 100 % | *L. amnigena* 98.42 % |
| TZW16 | GCA_016770955 | *L. amnigena* | *L. amnigena* 100 % | *L. amnigena* 98.42 % |
| UMA3121 | GCA_013337605 | *L. amnigena* | *L. amnigena* 100 % | *L. amnigena* 98.44 % |
| ZB04 | GCA_001652505 | *L. amnigena* | *H. chinensis* 96 % | *H. chinensis* 99.76 % |

\*MAGs; full names ERR1430553\_bin.131\_CONCOCT\_v1.1\_MAG, ERR1430553-bin.48 and ERR5094855\_bin.4\_metaWRAP\_v1.3\_MAG.

rMLST was used to provide tentative identifications for the *Lelliottia* genome sequences. As can be seen in **Table 2**, of the 23 genomes identified by NCBI as *L. amnigena*, only 19 were identified as *L. amnigena* with 100 % support by PubMLST, with two of the MAGs (ERR1430553, ERR1430553) identified as *L. amnigena* with low support scores. Strain 4928STDY7071390 (accession GCA_902160115) was identified as *L. nimipressuralis* (93 % support), while strain ZB04 was identified as *Huaxiibacter chinensis* (96 % support). Notable was identification of the proposed type strain of “*L. steviae*” (Lin *et al*. 2022) as *Pseudoalteromonas arabiensis* (100 % support). *L. jeotgali* is an earlier heterotypic synonym of *L. aquatilis* (Wu and Zong 2019), so we would expect the genomes of these species to share high support scores.

ANI analysis was undertaken to confirm identities of genomes (not shown). Identities determined by rMLST were confirmed for all genomes, except for strain A167 (accession GCA_021498285). An ANI of <95 % with the genome of the type strain of *L. amnigena* suggests this strain represents a novel species of *Lelliottia* (Chun *et al*. 2018). The genome of *L. jeotgali* shared 98.86 % ANI with that of *L. aquatilis*. Sourmash is a rapid method for computing hash sketches from genomic DNA sequences, and comparing them to each other. A comparison for sourmash signatures generated for all strains supported our findings from rMLST and ANI analyses (**Fig. 1b**). The sourmash analysis also confirmed the affiliation of GFKo1 with *L. amnigena*.

The genomes (*n*=19) of *L. amnigena* identified by rMLST to be *L. amnigena* (100 % support) and sharing ANI of >95 % with the genome of the type strain of *L. amnigena* were included in a phylogenetic analysis with the genomes of the type strains of *L. aquatilis* and *L. nimipressuralis* (**Fig. 1c**). All isolate-derived genomes clustered with the type strain of *L. amnigena*, while the MAG-derived sequence ERR5094855 clustered with *L. aquatilis* and *L. nimipressuralis*. The phylogenetic analysis confirmed the affiliation of GFKo1 with *L. amnigena*.

### Carriage of *bla*_LAQ-1_-like genes by *L. amnigena*

RGI/CARD analysis (loose, strict and perfect matches with protein sequences) showed strain GFKo1’s genome encoded no AMR genes. A pairwise alignment of GFKo1’s genome with the reference allele sequence of *bla*_LAQ-1_ (Li *et al*. 2022) showed GFKo1 encoded this class C β-lactamase, sharing 99.3 % nucleotide and 99.5 % amino acid pairwise identity with the reference sequence (accession MZ497396). In agreement with Li *et al*. (2022) we found that *bla*_LAQ-1_ encoded by GFKo1 had the obligatory serine active site of the β-lactamase catalytic motif S-V-S-K (serine-valine-serine-lysine) at positions 83–86, the typical class C β-lactamase motif Y-A-N (tryptophan-alanine-asparagine) at positions 169– 171, D/E (a peptide segment containing two dicarboxylic amino acids) at positions 236–238 and the conserved triad K-T-G (lysine-threonine-glycine) at positions 334–336 (**Supplementary Fig. 1**).

It is important to note that Bakta had annotated the *bla*_LAQ_ gene on contig 81 of GFKo1’s genome (locus tag GFKo1_06635). Among its databases, Bakta uses the NCBI Antimicrobial Resistance Gene Finder (AMRFinderPlus) (Feldgarden *et al*. 2022) to annotate AMR-associated genes in microbial genomes. In addition to a *bla*_LAQ-1_-like gene, AMRFinderPlus predicted GFKo1 to encode *vat* (Vat family streptogramin A *O*-acetyltransferase; GFKo1_06890), *catA* (type A chloramphenicol *O*-acetyltransferase; GFKo1_12820) and *oqxB* (multidrug efflux RND transporter permease subunit OqxB; GFKo1_19950). Bakta also predicted GFKo1 to encode the following AMR-associated genes: multidrug efflux MATE transporter EmmdR (GFKo1_03505); multidrug efflux MFS transporter EmrD (GFKo1_03800); Bcr/CflA family efflux transporter (GFKo1_04835); MdtK family multidrug efflux MATE transporter (GFKo1_04850); MATE efflux family protein (GFKo1_06250); multidrug efflux pump accessory protein AcrZ (GFKo1_15865); macrolide-specific efflux protein MacA (GFKo1_16470); putative aminoglycoside efflux pump (GFKo1_16810); multidrug efflux pump subunit AcrB (GFKo1_17175); multidrug efflux RND transporter periplasmic adaptor subunit AcrA (GFKo1_17180); multidrug efflux transporter transcriptional repressor AcrR (GFKo1_17185).

A BLASTP search of the predicted proteins in each of the genomes listed in **Table 1** against the amino acid sequence (380 aa) of the Bla_LAQ-1_ reference sequence identified one hit in each genome that shared >70 % identity and 100 % coverage with MZ497396 (**Supplementary Table 1**). The ‘hit’ sequences were extracted from the Bakta annotation files (available as **Supplementary Material**) for the genomes and used to create a multiple sequence alignment with the AmpC reference sequences included in the original characterization of *bla*_LAQ-1_ (Li *et al*. 2022). A phylogenetic analysis (maximum likelihood) demonstrated all the *L. amnigena* sequences clustered together (**Fig. 2**), sharing pairwise identity values of 98.16–99.47 % with Bla_LAQ-1_ of P13 and 97.63–100 % with each other (**Supplementary Table 2**), and high bootstrap support (97 %). The sequence of strain A167 (accession GCA_021498285) formed a branch on its own (100 % bootstrap support), providing additional support that this strain represents a novel species of *Lelliottia* (93.42 % amino acid identity with P13’s Bla_LAQ-1_ sequence). The sequences derived from *H. chinensis* strains clustered together but apart from the *L. amnigena* sequences, as did those of *L. nimipressuralis*, and those of *L. aquatilis* and *L. jeotgali* (all with 100 % bootstrap support).

**Fig. 2.**
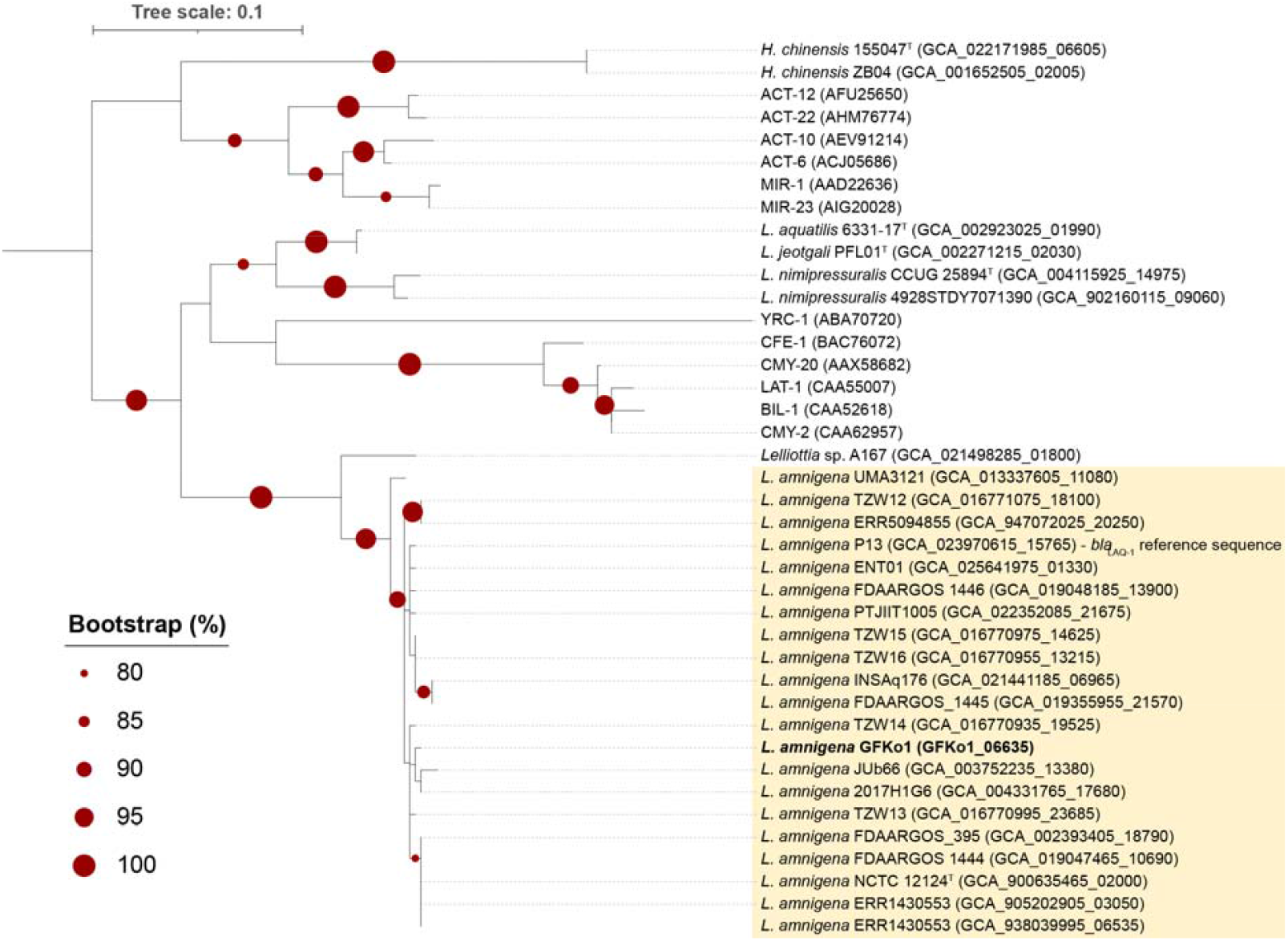
*bla*_LAQ_ is a core gene of *L. amnigena*. The *bla*_LAQ-1_ sequence of *L. amnigena* P13 represents the reference for this chromosomally-encoded AmpC β-lactamase (Li *et al*. 2022). Twelve other AmpC β-lactamases (ACT-12, ACT-22, BIL-1, CMY-2, CMY-20, LAT-1, CFE-1, YRC-1, MIR-1, MIR-23, ACT-6, ACT-10; (Li *et al*. 2022)) were included in the analysis for comparative purposes; the accessions for the amino acid sequences of these proteins are given in parentheses. The tree was rooted at the midpoint. Scale bar, average number of amino acid substitutions per position. The clade in yellow highlights *L. amnigena sensu stricto* sequences. Bootstrap values >80 % (based on 100 replications) are shown on the tree. The multiple sequence alignment used to create this phylogenetic tree is available as **Supplementary Material**.

### Phenotypic resistance profile of *L. amnigena* GFKo1

Disc diffusion assays were performed against antibiotics from a range of classes to determine the phenotypic resistance profile of *L. amnigena* GFKo1. Strain GFKo1 was found to be clinically sensitive to all antibiotics tested: penicillins (ampicillin, ampicillin-sulbactam, piperacillin, amoxicillin-clavulanate, piperacillin-tazobactam); cephalosporins (cefoxitin, ceftazidime, cefepime, cefotaxime, ceftriaxone); carbapenems (imipenem, meropenem, ertapenem); the monobactam aztreonam; the aminoglycosides amikacin and gentamicin; the fluoroquinolones ciprofloxacin and norfloxacin; the tetracyclines tigecycline and tetracycline; and trimethoprim and sulphamethoxazole-trimethoprim. A full table of results, including zone diameters measured and breakpoints can be found in **Supplementary Table 3**.

## DISCUSSION

In this study, we have characterized the genome and AMR genotype/phenotype of a strain of *L. amnigena* (GFKo1) isolated from the lung of a harbour porpoise stranded in 1993. We compared the genome of GFKo1 with genomes of closely related species (**Figure 1, Table 1** and **Table 2**), and demonstrated that *bla*_LAQ_, a chromosomally-encoded AmpC β-lactamase conferring resistance to penicillin G, ampicillin and several cephalosporins (Li *et al*. 2022), is a core gene of *L. amnigena* (**Figure 2**). Phenotypically, GFKo1 was sensitive to all antibiotics it was tested against, including ampicillin, cefotaxime and ceftazidime (**Supplementary Table 3**).

Our detailed genome-based identification of *L. amnigena* genomes (*n*=20 isolates; *n*=3 MAGs) downloaded from GenBank highlighted misclassification problems with four of the genomes, including that of a proposed type strain for “*L. steviae*” (Lin *et al*. 2022) (**Figure 1, Table 2**). While NCBI classifies some genome assemblies as anomalous and excludes them from the RefSeq database based on a range of different criteria, these assemblies are still available for download from GenBank. *Lelliottia* spp. data within NCBI GenBank are derived from isolates and MAGs, with no information provided as to, for example, the completeness and contamination of the genomes compared with accepted standards (Bowers *et al*. 2017). We have previously encountered problems with taxonomic assignments provided by NCBI (though acknowledge annotations are improving and being updated constantly; (Chen *et al*. 2020)). However, we still recommend that, for informative and accurate comparative genomic analyses to be undertaken, it is important that the genomes of all bacteria retrieved from public repositories are carefully checked for quality and identity before undertaking in-depth analyses.

In addition to identifying *bla*_LAQ_ as a core gene of *L. amnigena*, we demonstrated that proteins sharing high identity with a range of other AmpC β-lactamases were identified across all genomes included in this study (**Figure 2**). Whether these AmpC β-lactamases detected in non-*L. amnigena* genomes are functional remains to be determined. With respect to the *bla*_LAQ_ gene of GFKo1, it possessed the canonical motifs and active sites associated with β-lactamase enzymes. Additionally, it shared 99.5 % amino acid pairwise identity with LAQ-1 from *L. amnigena* P13 (accession MZ497396). It has been suggested that LAQ-1 from *L. amnigena* P13 confers resistance to a range of β-lactams, including first-to fourth-generation cephalosporins. A recombinant *Escherichia coli* clone of the β-lactamase from a plasmid-borne copy of *bla*_LAQ-1_ exhibited increased minimum inhibitory concentrations (MICs) to a range of antibiotics including ampicillin, cefoxitin, cefazolin, ceftazidime, cefepime, aztreonam, ticaracillin, piperacllin and cloxacillin. However, these increased MICs only resulted in clinical resistance to ampicillin, cefoxitin and cefazolin according to EUCAST guidelines. Despite the high level of sequence similarity between the *bla*_LAQ_ gene of GFKo1 and that from P13, *L. amnigena* GFKo1 was sensitive to all antibiotics tested in our study. Genomic alignment of the two strains showed a high level of sequence similarity in the region immediately upstream of the *bla*_LAQ-1_ gene, suggesting that lack of activity is not due to a mutation(s) in the promoter region.

Despite *bla*_LAQ_ being a core gene of all sequenced *L. amnigena* isolates, it is evident that broad-spectrum resistance to β-lactam antibiotics is not a uniform feature of the species. Resistance to penicillins is reported frequently, however resistance to specific cephalosporins is highly variable (Bollet *et al*. 1991; Stock and Wiedemann 2002; Murugaiyan *et al*. 2015; Li *et al*. 2022). Genome sequence data are rarely available for the strains characterised in these studies, making it impossible to determine the genotypic factors that contribute to the observed resistant phenotypes.

In summary, we show that the chromosomally-encoded AmpC β-lactamase *bla*_LAQ_ is a core gene of *L. amnigena*. However, presence of the *bla*_LAQ_ gene does not always correlate with phenotypic resistance to β-lactam antibiotics. Resistance to specific cephalosporins appears to be highly variable across the species. The mechanisms controlling *bla*_LAQ_ expression, and the degree to which *bla*_LAQ_ contributes to phenotypic resistance, require further investigation. Studies involving the cloning and expression of diverse *bla*_LAQ_ genes in genetic backgrounds free from other resistance markers will help elucidate the specificity of these novel β-lactamases and their role in *L. amnigena*.

## Abbreviations

AMR: antimicrobial resistance
ANI: average nucleotide identity
EUCAST: EUropean Committee on Antimicrobial Susceptibility Testing
MAG: metagenome-assembled genome
rMLST: ribosomal multi-locus sequence typing.

## ACKNOWLEDGEMENTS

GF isolated the strain and performed phenotypic characterization and antibiotic sensitivity test work. DN did all library preparation, sequencing and antibiotic sensitivity testing. LH did all bioinformatics work. DN and LH undertook all data analyses and interpretation. All authors contributed to the writing of the manuscript, and approved the submitted version.

---

**Supplementary Fig 1.**
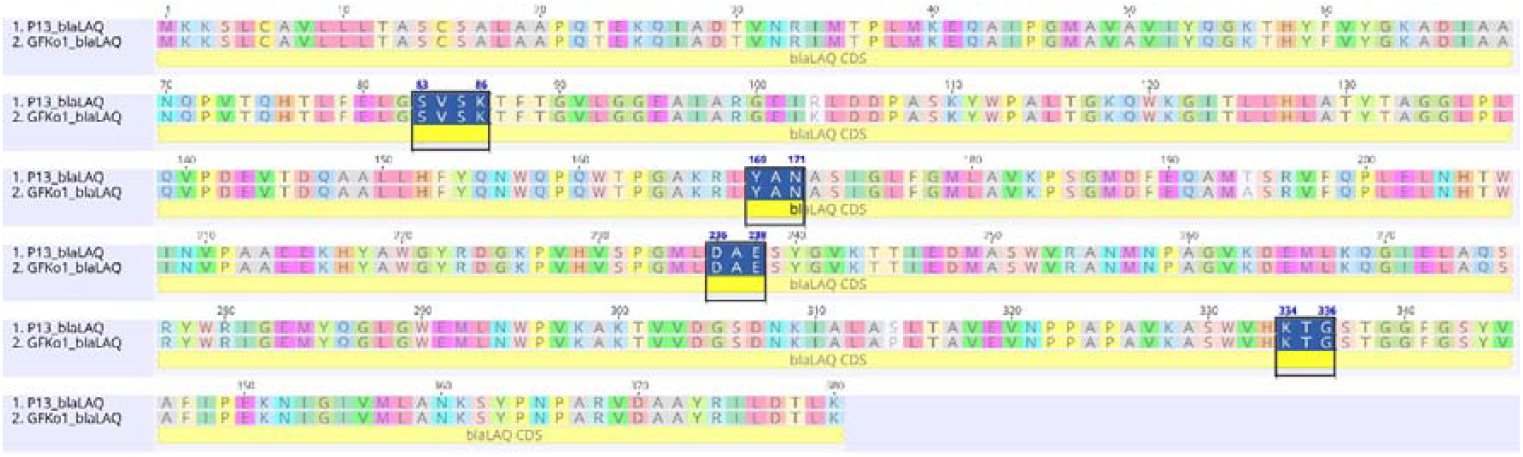
Amino acid alignment of β-lactamase LAQ-1 from *L. amnigena* P13 (Accession QXM27670) with β-lactamase LAQ-1 from *L. amnigena* GFko1 (locus tag GFKo1_06635). Conserved amino acid sequences typical for a class C β-lactamase are highlighted in blue.

**Supplementary Fig 2.**
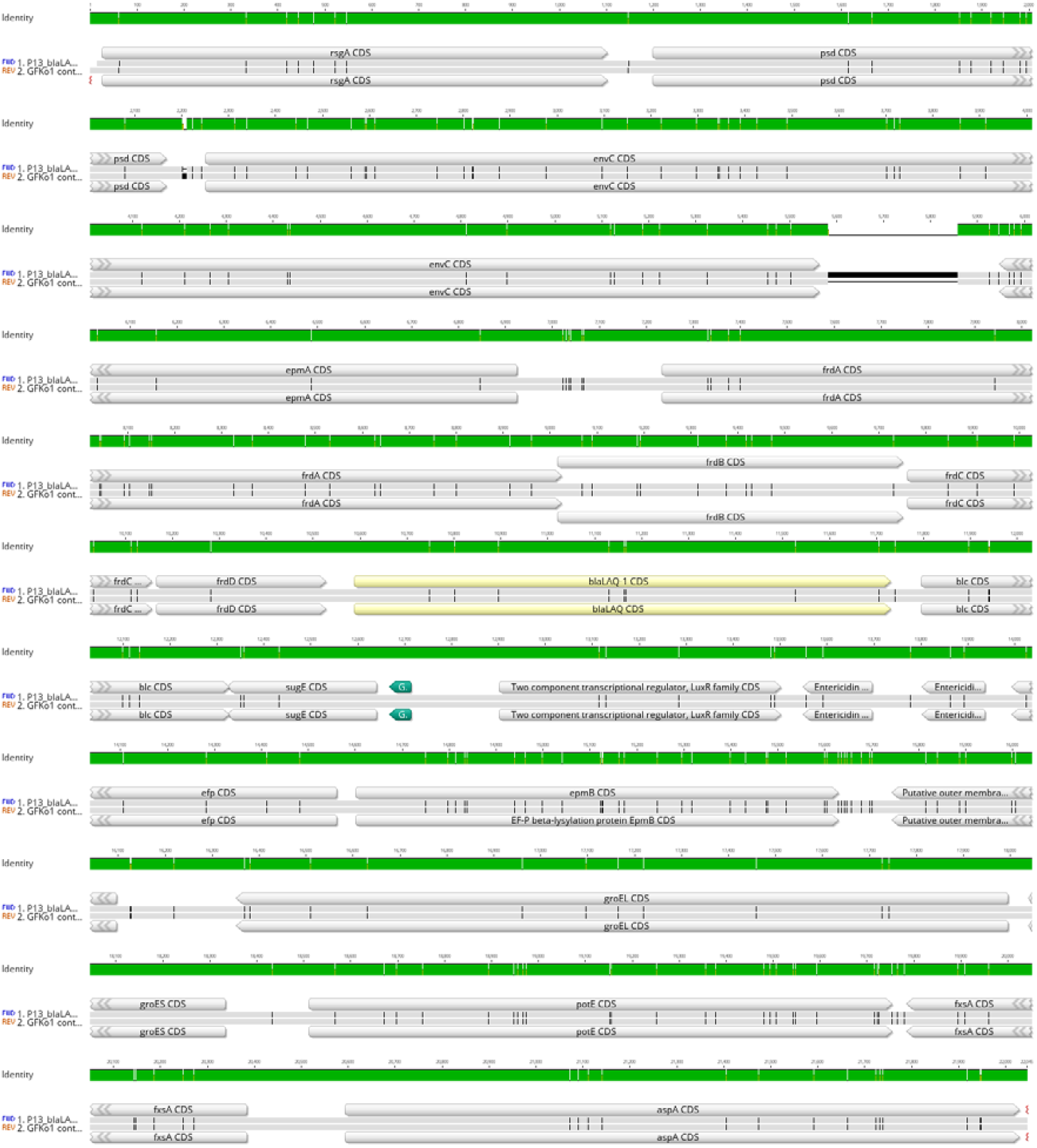
Alignment of *bla*_LAQ-1_-like and surrounding genome region of strain GFKo1 with the same region of *L. amnigena* P13. Gene predictions and annotations were made using Bakta as described in Methods. The sequence of GFKo1 was reverse-complemented and the MAFFT alignment shown was created in Geneious Prime v2023.0.1. The alignment covers 21,897 bp; the sequences share 21,497 identical sites (97.6 % pairwise identity) at the nucleotide level. The region shown matches that analysed by (Li *et al*. 2022); their analysis included sequence data from *L. amnigena* strains P13, NCTC 12124^T^, FDAARGOS 1444, FDAARGOS 1446, FDAARGOS_1445 and FDAARGOS_395.

